# Stratified Immune Profiling Uncovers Prognostic Heterogeneity Beyond MYCN Amplification and Age in Neuroblastoma

**DOI:** 10.64898/2026.07.15.738570

**Authors:** Jessica M. Magno, João C. D. Muzzi, Jean S. S. Resende, Lana B. P. Querne, Larissa M. Alvarenga, Luciane R. Cavalli, Bonald C. Figueiredo, Mauro A. A. Castro

## Abstract

Neuroblastoma is the most common extracranial solid tumor in children, presenting remarkable clinical heterogeneity with survival outcomes ranging from spontaneous regression to aggressive progression. MYCN oncogene amplification and age at diagnosis are established prognostic factors that are typically treated as independent covariates in risk stratification, yet their joint influence on the tumor immune microenvironment remains poorly understood. Here we show that stratifying patients by both variables simultaneously reveals six reproducible immune subtypes with distinct transcriptional programs and prognostic significance. Consensus clustering of immunomodulatory gene expression profiles from 149 patients in the TARGET-NBL cohort identified subtypes whose survival trajectories differ significantly within clinical strata defined by MYCN status and age at diagnosis. A linear Support Vector Machine classifier trained on these subtypes, using immunomodulatory gene expression combined with MYCN amplification status and age at diagnosis as predictive features, achieved 97.2% accuracy and a Cohen’s Kappa of 0.963 under 10-fold cross-validation, and generalized to an independent cohort of 493 patients (GSE62564). Kaplan-Meier analysis revealed significant survival differences across subtypes in both cohorts (TARGET-NBL: log-rank p = 0.0018; GSE62564: log-rank p < 0.0001). Single-sample gene set enrichment analysis identified differential activation of proliferative and immune response pathways across subtypes, consistent between both cohorts. These findings suggest that integrating MYCN amplification status and age at diagnosis as joint determinants of immune organization may reveal prognostic heterogeneity that is not fully captured when these factors are considered independently.

## 1 Introduction

Neuroblastoma is the most common extracranial solid tumor in childhood, accounting for 8–10% of all childhood malignancies and representing approximately 15% of all pediatric oncology deaths [1, 23]. Originating from neural crest progenitor cells, it can arise anywhere along the sympathetic nervous system, with the majority of primary tumors occurring in the abdomen [1, 7, 20]. The disease exhibits remarkable clinical heterogeneity, ranging from spontaneous regression in infants to aggressive metastatic progression resistant to intensive multimodal therapy [7, 23].

MYCN oncogene amplification is the strongest independent adverse prognostic biomarker in neuroblastoma, present in approximately 20% of all cases and contributing to roughly half of high-risk disease [7, 18]. Its identification automatically classifies the patient as high-risk regardless of age or disease stage, and it is frequently associated with unfavorable histology, segmental chromosomal alterations, and dysregulation of telomerase activity [7, 23]. Beyond its role as a prognostic marker, MYCN drives transcriptional programs that promote cell proliferation and suppress differentiation. These transcriptional effects extend to immune-related gene networks: emerging evidence indicates that MYCN actively shapes an immunosuppressive tumor microenvironment by repressing CXCL10-mediated recruitment of CD8^+^ T cells, downregulating CCL2-dependent natural killer cell chemotaxis, and reducing interferon pathway activity [18].

Age at diagnosis is among the most critical determinants of survival in neuroblastoma and is central to current risk stratification frameworks [23, 7]. The prognostic cutoff of greatest biological and clinical relevance lies at 18 months: patients diagnosed earlier have substantially better outcomes, including a higher probability of spontaneous regression in localized and metastatic 4S disease, whereas those diagnosed after 18 months present disproportionately with advanced, treatment-refractory metastatic disease [1, 23]. Current risk stratification integrates MYCN amplification status, age, histological classification, segmental chromosomal alterations, and ploidy [7], typically treating these variables as independent covariates rather than as joint determinants of tumor biology [12]. Whether a tumor regresses or progresses is determined by a combination of these factors [21], yet their intersection with immune microenvironment organization remains underexplored.

The tumor microenvironment in neuroblastoma comprises variable proportions of T cells, B cells, natural killer cells, and macrophages, with composition and functional states exhibiting complex associations with patient outcomes [13]. Intra- and inter-tumoral heterogeneity extends beyond tumor cell features to encompass the immune and non-immune cell populations of the TME, representing a major axis of disease complexity [13]. Systematic characterization of the immune landscape across human cancers by Thorsson et al. identified a pan-cancer set of immunomodulatory genes whose expression patterns reflect key functional states of antitumor immunity [25], providing a biologically grounded framework for immune microenvironment characterization that may be extended to tumor types not represented in the original cohort. Yet how MYCN amplification and age at diagnosis jointly determine immune microenvironment organization in neuroblastoma remains unexamined, a gap that limits the characterization of immune heterogeneity specific to their intersection [18, 12].

In this study, we explored this gap by integrating MYCN amplification status and age at diagnosis as joint determinants of immune microenvironment organization in neuroblastoma. Applying consensus clustering to immunomodulatory gene expression profiles in the TARGET-NBL cohort (n = 149), we identify six immune subtypes with distinct transcriptional programs and prognostic significance. A Support Vector Machine classifier trained on these subtypes generalizes to an independent validation cohort of 493 patients (GSE62564) from a distinct geographic and ethnic background, suggesting that the identified immune organization reflects consistent transcriptional properties of neuroblastoma rather than a cohort-specific artifact. Single-sample gene set enrichment analysis reveals differential activation of proliferative, immune, and developmental pathways across subtypes, providing a transcriptional basis for their distinct survival trajectories and suggesting that immune microenvironment organization within clinical strata may represent an additional dimension of prognostic heterogeneity in neuroblastoma.

## 2 Methods

### 2.1 Dataset and Cohorts

Gene expression and clinical data for the discovery cohort were obtained from the Therapeutically Applicable Research to Generate Effective Treatments Neuroblastoma project (TARGET-NBL, n = 149) [17], retrieved from the Genomic Data Commons (GDC) portal. RNA sequencing data were processed using variance-stabilizing transformation (VST) as implemented in DESeq2 [14]. Clinical annotations included MYCN amplification status, age at diagnosis, COG risk group, and INSS stage. For independent validation, the publicly available dataset GSE62564 (n = 498) was retrieved from the Gene Expression Omnibus (GEO) repository [2], comprising RNA-seq expression profiles (log2RPM-normalized) from 498 primary neuroblastoma tumor samples generated by the Sequencing Quality Control (SEQC) project [22, 26]. Clinical annotations, including MYCN amplification status and age at diagnosis, were obtained from the associated series GSE49711 [22]. Samples lacking MYCN amplification status information were excluded, resulting in a final validation cohort of 493 samples.

### 2.2 Immunomodulatory Gene Expression Analysis

A panel of immunomodulatory genes curated by Thorsson et al. [25] was used to characterize the tumor immune microenvironment. Of the 75 genes in the original panel, 71 were present in the TARGET-NBL expression matrix and retained for analysis. Genes were organized into seven functional categories: 1) co-stimulator, 2) co-inhibitor, 3) ligand, 4) receptor, 5) cell adhesion, 6) antigen presentation, and 7) others. Expression values were subset from the VST-normalized matrix and scaled per sample (z-score).

### 2.3 Immune Subtype Identification

Patients were stratified into four clinical groups by crossing MYCN amplification status (Amplified, Not Amplified) with age at diagnosis (Group A: ≤ 18 months; Group B: *>*18 months). The group comprising MYCN-amplified patients aged ≤ 18 months contained only three samples and was therefore excluded from clustering and designated as “unclassified”. Consensus clustering was independently applied to the remaining three groups using the ConsensusClusterPlus R package [28] with the following parameters: 1,000 resampling iterations, 80% item subsampling, 100% feature subsampling, hierarchical clustering with Euclidean distance. The optimal number of clusters (*k*) for each group was determined by jointly evaluating the consensus CDF plots and the relative change in area under the CDF curve (delta area). This procedure yielded six immune subtypes: subtypes 1 and 2 (MYCN-amplified, Group B); subtypes 1a and 2a (Not Amplified, Group A); and subtypes 1b and 2b (Not Amplified, Group B). Heatmaps were generated using the ComplexHeatmap R package [6]. To assess the transcriptional placement of the three unclassified patients, a sensitivity analysis was performed in which the trained SVM classifier was applied to their immunomodulatory gene expression profiles combined with their clinical variables, following the same feature construction procedure used for the remaining samples. This analysis assigned predicted subtype labels to all three patients without modifying the clustering solution.

### 2.4 SVM Classifier Training and Validation

A linear Support Vector Machine (SVM) classifier was trained using the `caret` R package [11] with 10-fold cross-validation on the TARGET-NBL discovery cohort (n = 146, excluding the three unclassified samples). The feature matrix combined z-score normalized expression values of the 71 immunomodulatory genes with MYCN amplification status and age group encoded as binary dummy variables, reflecting the clinical variables used to define the subtypes. Cross-validation predictions were saved across all folds to enable per-class performance evaluation using overall accuracy, Cohen’s Kappa, and per-class sensitivity, specificity, precision, and F1-score. The trained classifier was subsequently applied to the GSE62564 validation cohort (n = 493) using an identical feature matrix, assigning predicted immune subtype labels to all validation samples.

To assess the transcriptomic separation of immune subtypes, principal component analysis (PCA) and Uniform Manifold Approximation and Projection (UMAP) were applied to the z-score normalized expression matrix of the 71 immunomodulatory genes using the `prcomp` base function and the `umap` R package [16], respectively. UMAP was configured with `n_neighbors` = 12, `min_dist` = 0.3, Euclidean distance metric, 500 epochs, and a fixed random seed (42) for reproducibility. Analyses were restricted to the 146 classified samples. In the UMAP visualization, confidence ellipses were drawn at the 68% level (corresponding to one standard deviation under a bivariate normal assumption) to represent the central dispersion of each immune subtype.

### 2.5 Survival Analysis

Overall survival (OS) was assessed using Kaplan-Meier estimators fitted separately for each immune subtype. Statistical differences among survival curves were evaluated by the log-rank test. Cox proportional hazards regression was performed with immune subtype as the sole predictor variable, using subtype 2a as the reference category, as this group exhibited the most favorable survival profile. Since immune subtypes were defined by the combination of MYCN amplification status and age at diagnosis, these variables are collinear with subtype assignment and cannot be jointly included in a multivariable model. In the TARGET-NBL cohort, subtype 1a was excluded from the Cox model due to complete separation, as no mortality events were observed within this group during the follow-up period; this subtype was retained in the GSE62564 model, where events were observed. Proportionality of hazards was verified using the Schoenfeld residuals test. Analyses were performed using the `survival` and `survminer` R packages [24, 9] and applied independently to the TARGET-NBL discovery cohort and the GSE62564 validation cohort.

### 2.6 Pathway Enrichment Analysis

Pathway activity was characterized using single-sample gene set enrichment analysis (ssGSEA) based on the 50 Hallmark gene sets from the Molecular Signatures Database (MSigDB), retrieved via the `msigdbr` R package [4]. Gene set enrichment was computed per sample using the `fgseaMultilevel` function from the `fgsea` package [10], with expression values centered per gene relative to the cohort mean prior to analysis. Normalized enrichment scores (NES) were computed for each sample and visualized as heatmaps with Hallmark gene sets organized into eight functional categories: Immune, Proliferation, Metabolic, Signaling, Pathway, DNA damage, Cellular component, and Development. Analyses were performed independently for both the TARGET-NBL and GSE62564 cohorts. Differences in pathway enrichment across immune subtypes were characterized descriptively based on mean NES values per subtype, as ssGSEA was applied as a per-sample characterization tool rather than a hypothesis-testing framework. No formal statistical comparisons were performed, pathway-level multiple testing correction is not applicable, and results are intended as hypothesis-generating observations.

### 2.7 Cellular Deconvolution

To estimate the abundance of immune cell populations within each immune subtype, cellular deconvolution was performed using the EPIC algorithm [19] as implemented in the IOBR R package [29]. For the TARGET-NBL cohort, raw count data were converted to TPM following conversion of Ensembl gene identifiers to gene symbols. For the GSE62564 cohort, log2RPM-normalized expression values were reverted to linear scale prior to deconvolution. EPIC estimates the fractions of six immune and stromal cell types — B cells, CD4^+^ T cells, CD8^+^ T cells, NK cells, macrophages, and endothelial cells — as well as an uncharacterized cell fraction. Deconvolution was performed independently for each cohort. Differences in estimated cell fractions across immune subtypes were evaluated using the Kruskal-Wallis test, followed by pairwise Wilcoxon tests with false discovery rate correction, restricted to paired subtypes within the same clinical stratum. Analyses were performed using the `rstatix` R package [8].

## 3 Results

### 3.1 Immunomodulatory gene expression reveals immune heterogeneity in neuroblastoma

To characterize the immune landscape of neuroblastoma, we profiled the expression of 71 immunomodulatory genes curated by Thorsson et al. [25] across 149 patients from the TARGET-NBL cohort, organized into seven functional categories: co-stimulator, co-inhibitor, ligand, receptor, cell adhesion, antigen presentation, and others. Expression values were z-score normalized and, to preserve the clinical structure defined by MYCN amplification status and age at diagnosis, hierarchical clustering was applied independently within each predefined clinical stratum for visualization purposes, using Euclidean distance and Ward linkage. The resulting heatmap revealed distinct expression patterns associated with both clinical variables (Supplementary Fig. S1). MYCN-non-amplified tumors from patients aged 18 months or older showed higher overall expression of immunomodulatory genes, particularly in the receptor and antigen presentation categories. Among MYCN-amplified tumors, two broad expression profiles were apparent: one with uniformly low expression across all functional categories and another with elevated expression in multiple categories. Within the MYCN-non-amplified group, expression patterns differed according to age at diagnosis, with younger patients showing a distinct transcriptional profile relative to older patients. These observations motivated the application of consensus clustering independently within each clinical stratum, as described below and outlined in the analytical pipeline (Supplementary Fig. S2).

### 3.2 Consensus clustering defines six immune subtypes across clinical strata

Patients were stratified into four clinical groups by crossing MYCN amplification status with age at diagnosis. The group comprising MYCN-amplified patients aged 18 months or younger contained only three samples and was designated “unclassified” due to insufficient size for stable clustering. Consensus clustering was applied independently to the remaining three groups using ConsensusClusterPlus [28], with *k* = 2 selected in each group based on the delta area criterion (Supplementary Fig. S3). In all three groups, solutions with *k* ≥ 3 produced only marginal gains in the delta area and consistently isolated one or two borderline samples, suggesting that additional clusters did not improve solution stability. This procedure yielded six immune subtypes: subtypes 1 (n = 12) and 2 (n = 15) from MYCN-amplified patients older than 18 months; subtypes 1a (n = 6) and 2a (n = 24) from MYCN-non-amplified patients aged 18 months or younger; and subtypes 1b (n = 35) and 2b (n = 54) from MYCN-non-amplified patients older than 18 months (Fig. 1). Subtypes 1, 1a and 2b showed higher overall immunomodulatory gene expression relative to their paired subtypes within the same clinical stratum, particularly in antigen presentation, co-inhibitory receptor, and immune ligand categories, including CXCL9, CXCL10, and IFNG. Within each stratum, the two subtypes differed markedly in expression profile, indicating that immune heterogeneity exists independently of MYCN amplification status and age at diagnosis. A sensitivity analysis applying the trained SVM classifier to the three unclassified patients assigned two to subtype 1 and one to subtype 2a, suggesting that their transcriptional profiles are most consistent with subtypes identified in other strata; these results should be interpreted with caution given the very small sample size.

**Fig. 1.**
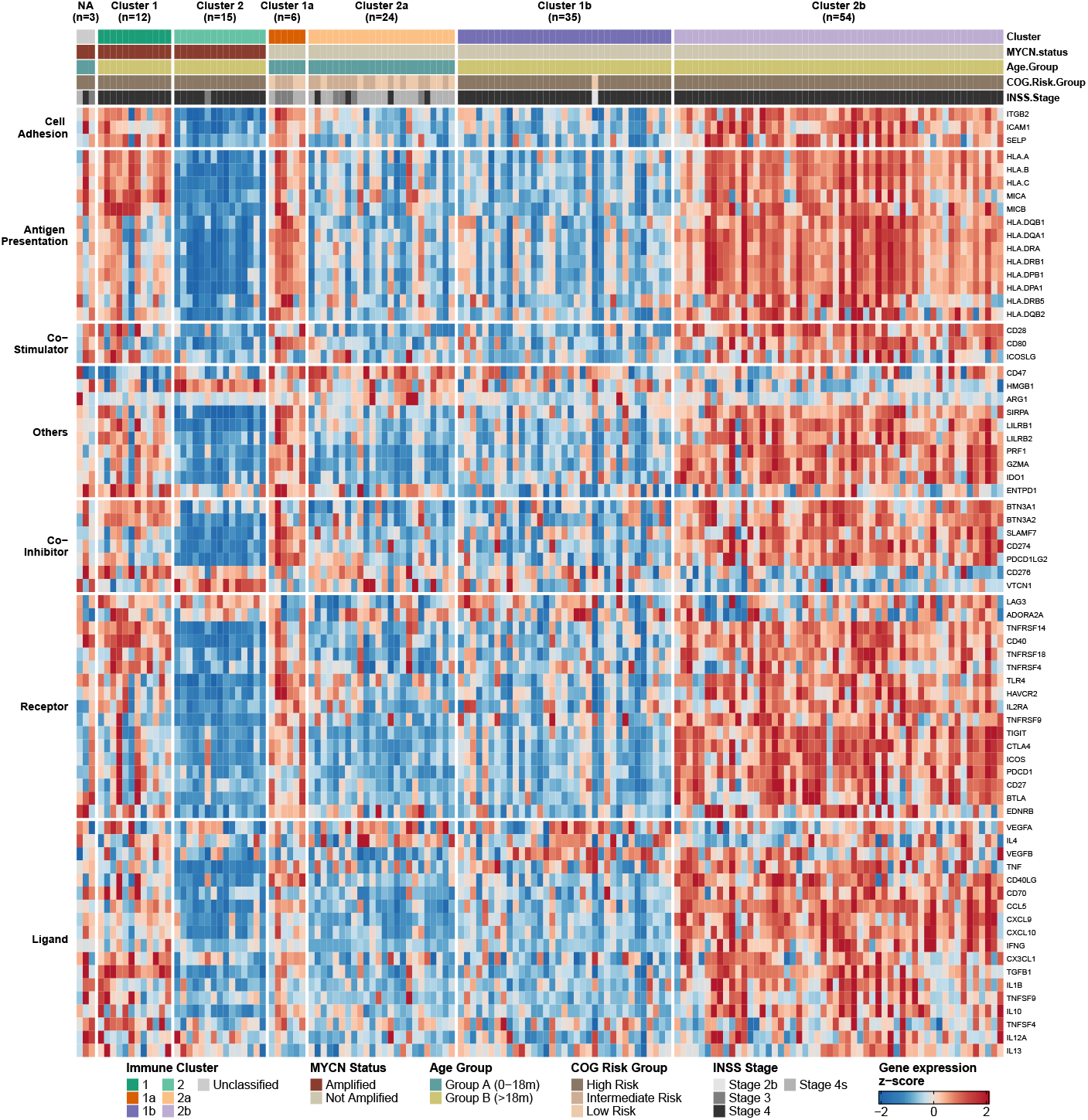
Six immune subtypes identified by consensus clustering in the TARGET-NBL cohort. Heatmap of z-score normalized expression of 71 immunomodulatory genes across 146 classified patients, ordered by immune subtype. The three Unclassified patients (MYCN-amplified, ≤ 18 months, n = 3) are shown separately. Gene rows are organized by functional category.

### 3.3 Immune subtypes occupy distinct transcriptomic spaces

To assess the transcriptomic separability of the six immune subtypes, we applied PCA and UMAP to the z-score normalized expression matrix of the 71 immunomodulatory genes. PCA was performed independently within each clinical stratum, consistent with the stratified clustering approach. Within the MYCN-amplified stratum, subtypes 1 and 2 showed clear separation along PC1, which captured 58.3% of variance (Supplementary Fig. S4A). Within the MYCN-non-amplified strata, separation was more gradual, with PC1 capturing 42.7% and 47.5% of variance in the younger and older age groups, respectively (Supplementary Fig. S4B–C), consistent with the broader transcriptional heterogeneity expected in larger and more clinically diverse groups. UMAP confirmed a structured organization of the six subtypes in the embedding space (Fig. 2A). Subtype 2 formed the most compact and isolated cluster, while subtypes 1b and 2a, derived from MYCN-non-amplified patients of different age groups, occupied partially overlapping regions, indicating shared immunomodulatory features across distinct age strata within the same MYCN background. Subtype 2b showed the broadest dispersion, consistent with greater transcriptional heterogeneity within the largest group (n = 54).

**Fig. 2.**
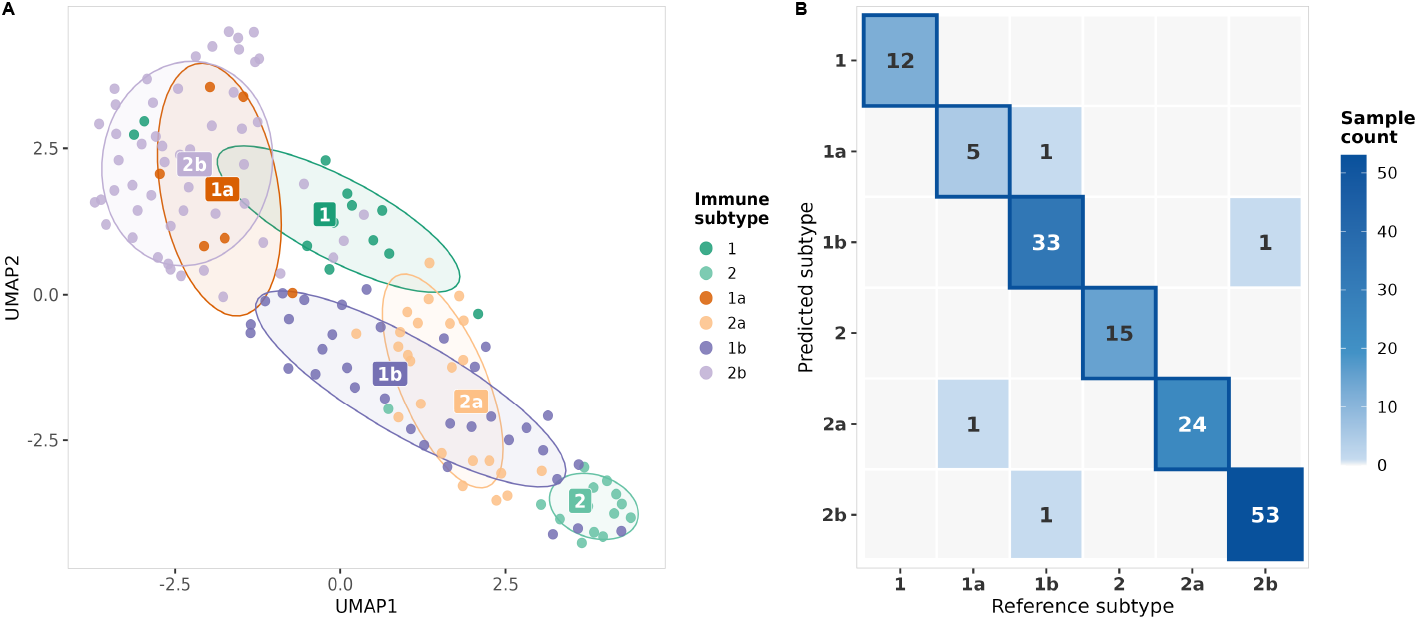
Transcriptomic separability and cross-cohort generalization of immune subtypes. **(A)** UMAP embedding of z-score normalized expression profiles from the TARGET-NBL cohort (n = 146), colored by immune subtype. Ellipses represent 68% confidence regions per subtype. **(B)** Cross-validated confusion matrix of the SVM classifier (10-fold CV, n = 146). Overall accuracy: 97.2%; Cohen’s Kappa: 0.963.

### 3.4 A linear SVM classifier generalizes immune subtype labels to an independent cohort

A linear Support Vector Machine classifier trained on the TARGET-NBL cohort with 10-fold cross-validation achieved an average accuracy of 97.2% and a Cohen’s Kappa of 0.963 (Fig. 2B, Table 1). The feature matrix combined z-score normalized expression values of the 71 immunomodulatory genes with MYCN amplification status and age group encoded as binary dummy variables, and the three Unclassified samples were excluded from model training.

**Table 1.** Per-class performance metrics of the SVM classifier evaluated by 10-fold cross-validation on the TARGET-NBL cohort (n = 146).

| Subtype | Sensitivity | Specificity | Precision | F1-score |
| --- | --- | --- | --- | --- |
| 1 | 1.000 | 1.000 | 1.000 | 1.000 |
| 1a | 0.833 | 0.993 | 0.833 | 0.833 |
| 1b | 0.943 | 0.991 | 0.971 | 0.957 |
| 2 | 1.000 | 1.000 | 1.000 | 1.000 |
| 2a | 1.000 | 0.992 | 0.960 | 0.980 |
| 2b | 0.981 | 0.989 | 0.981 | 0.981 |
| Overall accuracy: 97.2%; Cohen’s Kappa: 0.964 |  |  |  |  |

The classifier was applied to the GSE62564 validation cohort (n = 493) using an identical feature matrix, assigning all six immune subtypes with predicted distributions of subtype 1 (n = 30), 2 (n = 59), 1a (n = 119), 2a (n = 114), 1b (n = 65), and 2b (n = 106). Heatmap visualization confirmed that expression patterns in the validation cohort were consistent with those observed in the discovery cohort. Within each clinical stratum, the contrasting expression profiles between paired subtypes were maintained: subtypes 1, 1a and 2b showed higher overall immunomodulatory gene expression, while subtypes 2, 2a, and 1b showed comparatively lower expression across most gene categories (Supplementary Fig. S5). The immune expression patterns associated with the combination of MYCN amplification status and age at diagnosis were reproducible across cohorts of distinct geographic and ethnic origin.

### 3.5 Immune subtypes stratify patient survival in both cohorts

Kaplan-Meier analysis revealed significant differences in overall survival across the six immune subtypes in both the TARGET-NBL discovery cohort (logrank p = 0.0018, Fig. 3A) and the GSE62564 validation cohort (log-rank p < 0.0001, Fig. 3B). In both cohorts, subtypes 1a and 2a, derived from MYCN-non-amplified patients aged 18 months or younger, showed the most favorable outcomes, with survival probabilities above 87% at 60 months. Subtypes 1b and 2b, derived from MYCN-non-amplified patients older than 18 months, showed substantially lower survival in both cohorts, with subtype 1b reaching approximately 33% at 60 months in the discovery cohort. This contrast between subtypes sharing the same MYCN background but differing in age at diagnosis indicates that age contributes independently to survival outcomes within the MYCN-non-amplified group.

**Fig. 3.**
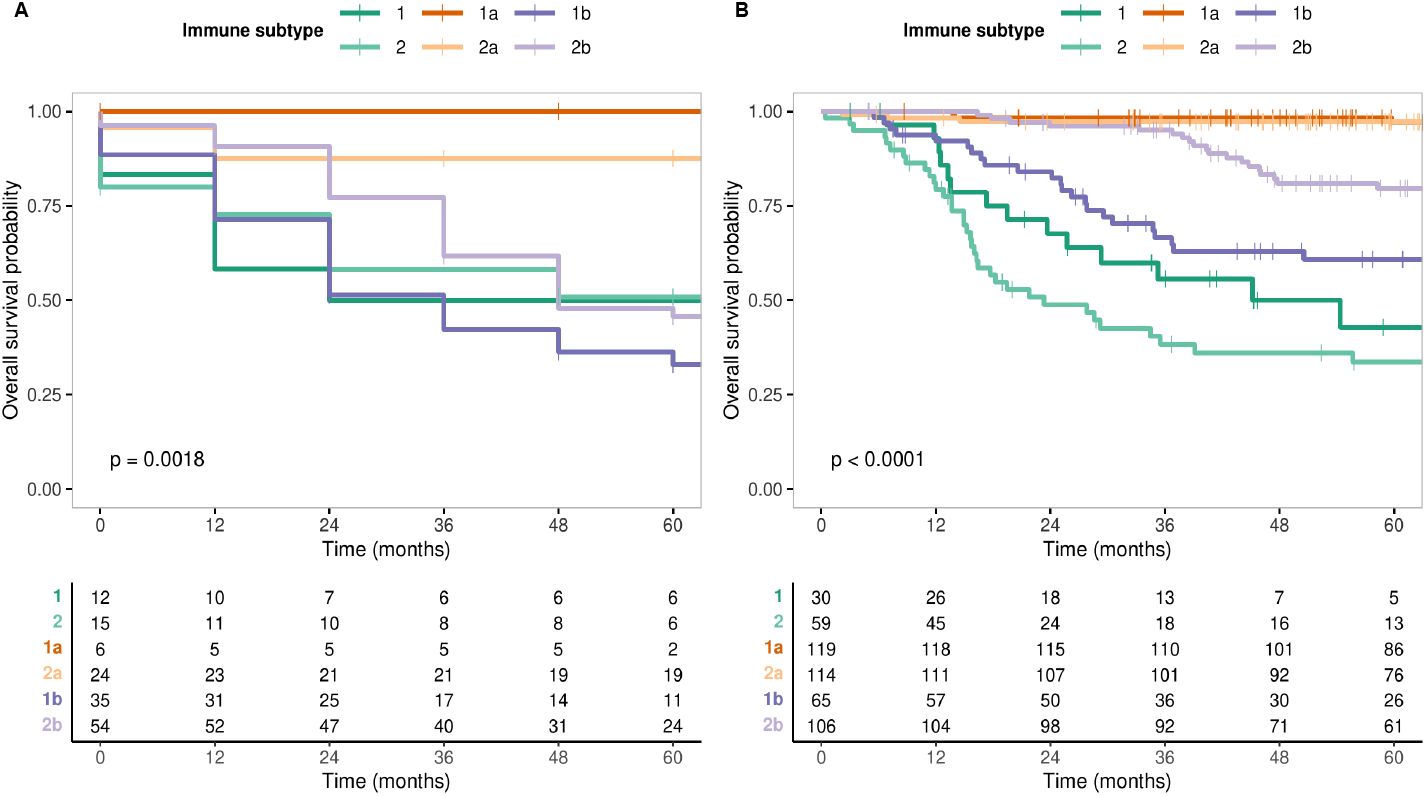
Kaplan-Meier overall survival curves by immune subtype. **(A)** TARGET-NBL discovery cohort (n = 146, log-rank p = 0.0018). **(B)** GSE62564 validation cohort (n = 493, log-rank p < 0.0001). Tick marks indicate censored observations.

Cox proportional hazards regression in the TARGET-NBL cohort was performed using subtype 2a as the reference category, as this group exhibited the most favorable survival profile. Subtype 1a was excluded from the model due to complete separation, as no mortality events were observed within this group. The model confirmed a significant global association between immune subtype and overall survival (log-rank p = 0.003, concordance index = 0.64, Supplementary Fig. S6A). Subtype 1b showed the highest mortality risk relative to subtype 2a (HR = 5.4, 95% CI: 2.05–14.1, p < 0.001), followed by subtype 2 (HR = 3.9, 95% CI: 1.27–11.9, p = 0.017), subtype 2b (HR = 3.1, 95% CI: 1.19–8.0, p = 0.020), and subtype 1 (HR = 3.2, 95% CI: 0.96–10.4, p = 0.058). The proportionality of hazards assumption was satisfied in the discovery cohort (Schoenfeld test p = 0.092).

In the GSE62564 validation cohort, Cox regression was performed including all six subtypes, as subtype 1a had sufficient events for estimation. Using sub-type 2a as the reference category, the model showed a strong global association between immune subtype and overall survival (log-rank p < 0.001, concordance index = 0.82, Supplementary Fig. S6B). Subtype 2 showed the highest mortality risk (HR = 34.3, 95% CI: 12.1–96.8, p < 0.001), followed by subtype 1 (HR = 23.3, 95% CI: 7.6–71.0, p < 0.001), subtype 1b (HR = 14.6, 95% CI: 5.1–42.0, p < 0.001), and subtype 2b (HR = 6.4, 95% CI: 2.2–18.6, p < 0.001). Subtype 1a did not differ significantly from subtype 2a (HR = 0.69, 95% CI: 0.15–3.1, p = 0.626), consistent with similarly favorable outcomes observed in the Kaplan-Meier analysis. However, the proportionality of hazards assumption was not satisfied in this cohort (Schoenfeld test p < 0.001). Given this violation, the hazard ratios reported for the GSE62564 cohort should be interpreted with caution, as they represent average effects over the entire follow-up period and may not accurately reflect the time-varying nature of the association between immune subtypes and survival.

### 3.6 Distinct pathway activation programs characterize each immune subtype

Single-sample gene set enrichment analysis of the 50 MSigDB Hallmark gene sets revealed distinct pathway activation patterns across the six immune subtypes, consistent between the TARGET-NBL and GSE62564 cohorts (Fig. 4). MYCN-amplified subtypes 1 and 2 showed enrichment of proliferative gene sets, including G2M checkpoint, E2F targets, and MYC targets, in both cohorts, with subtype 2 exhibiting the strongest enrichment of these signatures. In contrast, subtypes 1, 1a, and 2b showed enrichment of immune gene sets, including interferon alpha and gamma response, allograft rejection, and inflammatory response, consistently across both cohorts. These observations suggest that immune pathway activation patterns are not determined solely by MYCN amplification status or age at diagnosis, but may reflect a more complex transcriptional organization that emerges from the joint stratification by both clinical variables. These findings are descriptive and intended as hypothesis-generating, as no formal statistical comparisons between subtypes were performed.

**Fig. 4.**
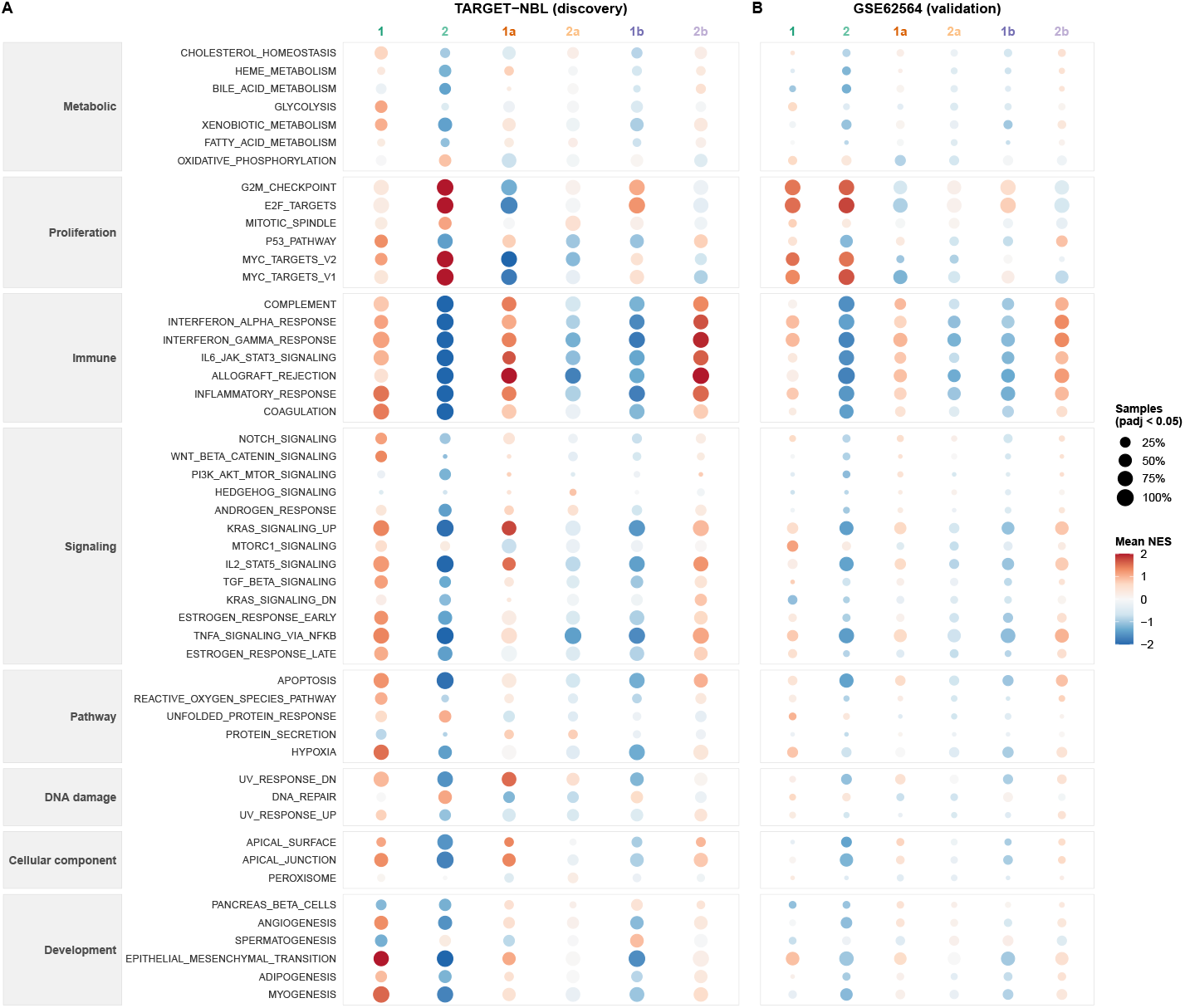
Pathway enrichment profiles across immune subtypes. Single-sample gene set enrichment analysis (ssGSEA) of 50 MSigDB Hallmark gene sets in **(A)** the TARGET-NBL discovery cohort and **(B)** the GSE62564 validation cohort. Dot color represents mean normalized enrichment score (NES) per subtype; dot size represents the fraction of samples with adjusted p < 0.05. Gene sets are ordered by functional category and mean NES across both cohorts, enabling direct visual comparison between panels. Functional categories are indicated in the left margin of panel A.

### 3.7 Cellular deconvolution supports immune heterogeneity across subtypes

To complement the transcriptional characterization of immune subtypes, cellular deconvolution using EPIC was applied to both cohorts. Estimated immune cell fractions were consistently low across all subtypes, consistent with the characteristically sparse immune infiltration of neuroblastoma. Nevertheless, statistically significant differences were observed for all cell types evaluated in both cohorts (Kruskal-Wallis, FDR-adjusted p<0.05; Supplementary Fig. S7). Macrophage and endothelial cell fractions showed the most consistent patterns across cohorts. Within each clinical stratum, estimated macrophage fractions were higher in subtypes 1 and 1a relative to their paired subtypes 2 and 2a, respectively, in both cohorts. Within the MYCN-non-amplified Group B stratum, subtype 2b showed higher estimated macrophage fractions than subtype 1b in both cohorts. A similar directional pattern was observed for endothelial cells, with subtypes 1 and 1a showing higher estimated fractions than subtypes 2 and 2a in both cohorts. Estimated CD4^+^ T cell fractions were significantly higher in subtype 1a versus 2a and in subtype 2b versus 1b in both cohorts, while NK cell fractions were near zero across all subtypes, consistent with the known paucity of NK cell infiltration in neuroblastoma. Collectively, these findings suggest that the immune heterogeneity identified at the transcriptional level is partially reflected in estimated immune cell composition, supporting the biological coherence of the identified subtypes. Given the inherent limitations of bulk RNA-seq deconvolution and the low absolute fractions observed, these results should be interpreted as hypothesis-generating estimates rather than direct measures of cellular infiltration.

## 4 Discussion

The stratification of neuroblastoma patients according to MYCN amplification status and age at diagnosis reveals that, even within these clinically defined groups, substantial variation exists in the immune transcriptional profile of the tumor microenvironment. Six reproducible immune subtypes, identified by consensus clustering applied within each clinical stratum, exhibit distinct associations with overall survival, indicating that the immune profile represents an additional layer of prognostic heterogeneity beyond that captured by established clinical risk stratification criteria [1, 21]. The reproducibility of these subtypes in an independent validation cohort of 493 patients from a distinct geographic and ethnic background suggests that this immune organization may reflect an intrinsic biological property of the tumor rather than a cohort-specific artifact. Collectively, these findings indicate that integrating immune transcriptional profiling into existing clinical classification may uncover prognostic heterogeneity that remains overlooked when MYCN amplification and age are considered independently of tumor immune composition.

Recent studies have characterized the immune landscape of neuroblastoma and its association with clinical outcomes, establishing that tumor immune composition carries prognostic information independent of established clinical variables such as age and MYCN amplification status [13]. Louault et al. further propose that combining genomic and tumor microenvironment information will be essential for improving risk stratification in high-risk neuroblastoma [13]. Supporting the biological rationale for this joint analysis, pan-neuroblastoma genomic profiling has shown that driver alterations, including MYCN amplification, differ markedly in frequency across age groups, indicating that age and molecular tumor features are not independent dimensions of disease biology [3]. However, in both the immune landscape characterizations and the genomic profiling studies, age and MYCN amplification are treated as covariates to adjust for rather than as joint determinants of immune organization, which limits the ability to detect immune heterogeneity specific to their intersection. By applying consensus clustering within strata defined by both variables simultaneously, the present study demonstrates that immune transcriptional variation with prognostic relevance exists within clinical risk groups, as evidenced by the distinct overall survival trajectories observed among subtypes sharing the same MYCN status and age stratum in both cohorts.

The transcriptional profiles of MYCN-amplified subtypes are consistent with mechanisms of MYCN-driven immunosuppression described by Qin et al. [18]. In subtype 2, ssGSEA revealed reduced enrichment of interferon alpha and gamma response, inflammatory response, and IL6/JAK/STAT3 signaling pathways in both cohorts, consistent with the suppression of CXCL10-mediated immune recruitment and reduced interferon pathway activity reported in MYCN-driven tumors, which has been associated with decreased CD8^+^ T cell infiltration at the cellular level [18]. Subtype 1, also MYCN-amplified, showed a contrasting profile with enrichment of immune response pathways, indicating that MYCN amplification alone does not uniformly determine immune pathway activation. Among MYCN-non-amplified subtypes, subtype 1b showed comparatively lower immune pathway enrichment relative to subtype 2b, suggesting that age-related factors may contribute independently to immune transcriptional variation through mechanisms other than MYCN-driven programs. This observation remains to be investigated experimentally and cannot be attributed to a specific molecular mechanism based on transcriptional data alone.

The transcriptional suppression signature observed in MYCN-non-amplified subtype 1b is consistent with findings reported by Masih et al., who identified distinct immune microenvironment subgroups within MYCN-non-amplified neuroblastoma using whole transcriptome sequencing [15]. In that study, a subset of MYCN-non-amplified tumors from older high-risk patients presented an immunosuppressive microenvironment characterized by stromal enrichment and immune cell infiltration patterns associated with worse clinical outcomes [15]. The convergence of these transcriptional observations with the immune subgroups described by Masih et al. suggests that the transcriptional profile of subtype 1b may reflect a similar immunosuppressive organization, though the underlying cellular composition cannot be inferred directly from bulk expression data and remains to be characterized experimentally. Cellular deconvolution using EPIC provided partial support for this interpretation, with estimated macrophage fractions consistently higher in subtypes associated with lower immune pathway enrichment within each clinical stratum in both cohorts. This pattern is consistent with the characterization of tumor-associated macrophages as a predominant immunosuppressive component of the neuroblastoma TME, where M2-polarized macrophages have been associated with disease progression and worse clinical outcomes [13, 15]. Estimated NK cell fractions were near zero across all subtypes, consistent with the known susceptibility of the neuroblastoma TME to NK cell exclusion and dysfunction driven by soluble factors such as TGF-*β* [13]. These estimates should be interpreted with caution given the inherent limitations of bulk RNA-seq deconvolution for resolving low-abundance cell populations in heterogeneous tumor samples [13, 15].

Cox proportional hazards analysis in the TARGET-NBL cohort confirmed that immune subtypes are independently associated with overall survival relative to the reference subtype 2a, with hazard ratios ranging from 3.1 to 5.4 for subtypes with poorer prognosis and a concordance index of 0.63, indicating moderate discriminative ability. Subtype 1, composed of MYCN-amplified patients, showed a trend toward increased hazard (HR 3.2; 95% CI 0.96–10; p = 0.058) that did not reach statistical significance, likely reflecting the limited sample size within that stratum. In the GSE62564 validation cohort, hazard ratios were larger in magnitude and more precisely estimated (concordance index 0.82), with MYCN-amplified subtypes showing increased risk relative to the reference (HR 23.29 and 34.27 for subtypes 1 and 2, respectively). However, the Schoenfeld residuals test indicated a violation of the proportional hazards assumption in this cohort (p = 0.00079) [5], suggesting that the association between immune subtypes and survival may vary over the follow-up period. This may reflect differences in follow-up duration, age composition, or treatment protocols between cohorts. Time-varying effects models could be explored in future analyses with more complete follow-up data.

This study has limitations that merit consideration. Immune subtype identification relies on bulk gene expression data, which captures average transcriptional signals across heterogeneous cell populations and does not allow direct quantification of specific immune cell types within the tumor microenvironment. Patient distribution across subtypes is unequal (n = 6 to 54 in TARGET-NBL), which may affect the stability of smaller subtypes and per-class classification metrics within a six-class design. The validation cohort shares the same sequencing platform and consortium of origin as the discovery cohort, limiting the assessment of full generalizability, and the absence of treatment annotation in GSE62564 precludes adjustment for therapeutic differences between cohorts; since treatment was not a study variable, this limitation pertains to the external interpretation of survival associations rather than to the validity of the analyses. The SVM classifier incorporates the same variables used to define the stratification structure as predictive features, which may in part inflate the reported classification performance. Future studies integrating single-cell RNA sequencing could provide more direct cellular characterization [27], and prospective validation in cohorts with complete clinical annotation would help clarify the prognostic relevance of these subtypes across different therapeutic contexts.

## 5 Conclusion

This study demonstrates that immune transcriptional profiles vary markedly within clinical strata defined by MYCN amplification status and age at diagnosis, and that this variation carries independent prognostic relevance in neuroblastoma. The six immune subtypes identified through consensus clustering exhibit distinct survival trajectories in both the discovery and validation cohorts, indicating that immune microenvironment organization represents a layer of biological heterogeneity not captured by current risk stratification frameworks. The cross-cohort reproducibility of these subtypes, supported by a machine learning classifier with high discriminative performance, suggests that the identified patterns reflect consistent transcriptional properties of neuroblastoma rather than cohort-specific phenomena. Integrating immune transcriptional and cellular deconvolution data into existing clinical classification may contribute to a more refined characterization of patient subgroups, with potential implications for future stratification strategies in neuroblastoma.

## Acknowledgments

This study was supported by the Instituto de Pesquisa Pelé Pequeno Príncipe (doctoral fellowship awarded to J.M.M.).

## Declaration of Generative AI and AI-Assisted Technologies in the Writing Process

During the preparation of this work, the authors used Claude (claude.ai, Anthropic) to improve text harmonization and manuscript readability. After using this tool, the authors carefully reviewed and edited all content as needed and take full responsibility for the content of this publication.

## References

1. Ahmed, A.A., Zhang, L., Reddivalla, N., Hetherington, M.: Neuroblastoma in children: Update on clinicopathologic and genetic prognostic factors. Pediatric Hematology and Oncology 34(3), 165–185 (2017). 10.1080/08880018.2017.1330375

2. Barrett, T., Wilhite, S.E., Ledoux, P., et al.: NCBI GEO: archive for functional genomics data sets – update. Nucleic Acids Research 41(D1), D991–D995 (2013). 10.1093/nar/gks1193

3. Brady, S.W., Liu, Y., Ma, X., Gout, A.M., et al.: Pan-neuroblastoma analysis reveals age- and signature-associated driver alterations. Nature Communications 11, 5183 (2020). 10.1038/s41467-020-18987-4

4. Dolgalev, I.: msigdbr: MSigDB gene sets for multiple organisms in a tidy data format (2022), https://CRAN.R-project.org/package=msigdbr, r package version 7.5.1

5. Grambsch, P.M., Therneau, T.M.: Proportional hazards tests and diagnostics based on weighted residuals. Biometrika 81(3), 515–526 (1994). 10.1093/biomet/81.3.515

6. Gu, Z., Eils, R., Schlesner, M.: Complex heatmaps reveal patterns and correlations in multidimensional genomic data. Bioinformatics 32(18), 2847–2849 (2016). 10.1093/bioinformatics/btw313

7. Irwin, M.S., Goldsmith, K.C.: Current and emerging biomarkers: Impact on risk stratification for neuroblastoma. Journal of the National Comprehensive Cancer Network 22(6), e247051 (2024). 10.6004/jnccn.2024.7051

8. Kassambara, A.: krstatix: Pipe-Friendly Framework for Basic Statistical Tests (2023), https://CRAN.R-project.org/package=rstatix, r package version 0.7.2

9. Kassambara, A., Kosinski, M., Biecek, P.: survminer: Drawing survival curves using ggplot2 (2021), https://CRAN.R-project.org/package=survminer, r package version 0.4.9

10. Korotkevich, G., Sukhov, V., Budin, N., Shpak, B., Artyomov, M.N., Sergushichev, A.: Fast gene set enrichment analysis. bioRxiv (2021). 10.1101/060012

11. Kuhn, M.: Building predictive models in R using the caret package. Journal of Statistical Software 28(5), 1–26 (2008). 10.18637/jss.v028.i05

12. Li, X., Li, W., Wang, J.: Integrative analysis of CRISPR screening and gene expression data identifies a three-gene prognostic model associated with immune microenvironment in neuroblastoma. Translational Cancer Research (2025). 10.21037/tcr-2024-2472

13. Louault, K., De Clerck, Y.A., Janoueix-Lerosey, I.: The neuroblastoma tumor microenvironment: From an in-depth characterization towards novel therapies. EJC Paediatric Oncology 3, 100161 (2024). 10.1016/j.ejcped.2024.100161

14. Love, M.I., Huber, W., Anders, S.: Moderated estimation of fold change and dispersion for RNA-seq data with DESeq2. Genome Biology 15, 550 (2014). 10.1186/s13059-014-0550-8

15. Masih, K.E., Wei, J.S., Milewski, D., Khan, J.: Exploring and targeting the tumor immune microenvironment of neuroblastoma. Journal of Cellular Immunology 3(5), 305–316 (2021). 10.33696/immunology.3.111

16. McInnes, L., Healy, J., Melville, J.: UMAP: Uniform manifold approximation and projection for dimension reduction. arXiv (2018). 10.48550/arXiv.1802.03426

17. Pugh, T.J., Morozova, O., Attiyeh, E.F., Asgharzadeh, S., et al.: The genetic landscape of high-risk neuroblastoma. Nature Genetics 45(3), 279–284 (2013). 10.1038/ng.2529

18. Qin, X., Lam, A., Zhang, X., Sengupta, S., et al.: CKLF instigates a “cold” microenvironment to promote MYCN-mediated tumor aggressiveness. Science Advances 10 (2024). 10.1126/sciadv.adh9547

19. Racle, J., de Jonge, K., Baumgaertner, P., Speiser, D.E., Gfeller, D.: Simultaneous enumeration of cancer and immune cell types from bulk tumor gene expression data. eLife 6, e26476 (2017). 10.7554/eLife.26476

20. Ratner, N., Brodeur, G.M., Dale, R.C., Schor, N.F.: The “neuro” of neuroblastoma: Neuroblastoma as a neurodevelopmental disorder. Annals of Neurology 80(1), 13–23 (2016). 10.1002/ana.24659

21. Rivera, Z., Escutia, C., Madonna, M.B., Gupta, K.H.: Biological insight and recent advancement in the treatment of neuroblastoma. International Journal of Molecular Sciences 24(10), 8470 (2023). 10.3390/ijms24108470

22. SEQC/MAQC-III Consortium: A comprehensive assessment of RNA-seq accuracy, reproducibility and information content by the Sequencing Quality Control Consortium. Nature Biotechnology 32(9), 903–914 (2014). 10.1038/nbt.2957

23. Takita, J.: Molecular basis and clinical features of neuroblastoma. JMA Journal 4(4), 321–331 (2021). 10.31662/jmaj.2021-0077

24. Therneau, T.M.: A Package for Survival Analysis in R (2024), https://CRAN.R-project.org/package=survival, r package version 3. 8–3

25. Thorsson, V., Gibbs, D.L., Brown, S.D., et al.: The immune landscape of cancer. Immunity 48(4), 812–830 (2018). 10.1016/j.immuni.2018.03.023

26. Wang, C., Gong, B., Bushel, P.R., Thierry-Mieg, J., et al.: The concordance between RNA-seq and microarray data depends on chemical treatment and transcript abundance. Nature Biotechnology 32(9), 926–932 (2014). 10.1038/nbt.3001

27. Wienke, J., Visser, L.L., et al.: Integrative analysis of neuroblastoma by single-cell RNA sequencing identifies the NECTIN2-TIGIT axis as a target for immunotherapy. Cancer Cell 42 (2024). 10.1016/j.ccell.2023.12.008

28. Wilkerson, M.D., Hayes, D.N.: ConsensusClusterPlus: a class discovery tool with confidence assessments and item tracking. Bioinformatics 26(12), 1572–1573 (2010). 10.1093/bioinformatics/btq170

29. Zeng, D., Ye, Z., Shen, R., Yu, G., Wu, J., et al.: IOBR: Multi-omics immuno-oncology biological research to decode tumor microenvironment and signatures. Front. Immunol. 12, 687975 (2021). 10.3389/fimmu.2021.687975

