## Supplementary Figures S1-S7 for "Stratified Immune Profiling Uncovers Prognostic Heterogeneity Beyond MYCN Amplification and Age in Neuroblastoma"

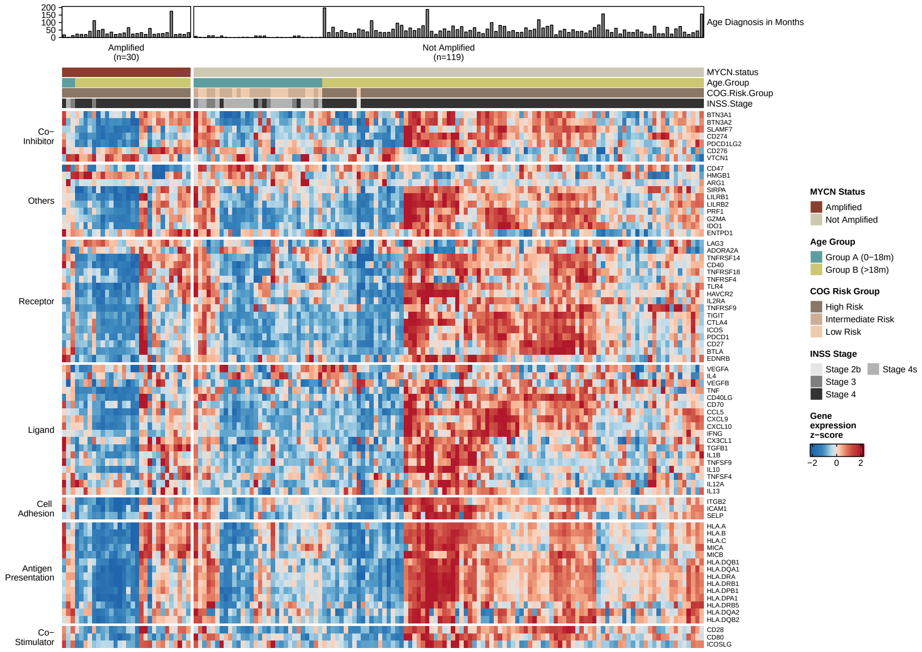


**Fig. S1.** Immunomodulatory gene expression in the TARGET-NBL cohort. Heatmap of z-score normalized expression (clamped to $\pm 2$) of 71 immunomodulatory genes across 149 patients, organized into seven functional categories. Samples are ordered by hierarchical clustering applied independently within each clinical stratum and annotated by MYCN amplification status, age group, COG risk group, and INSS stage.

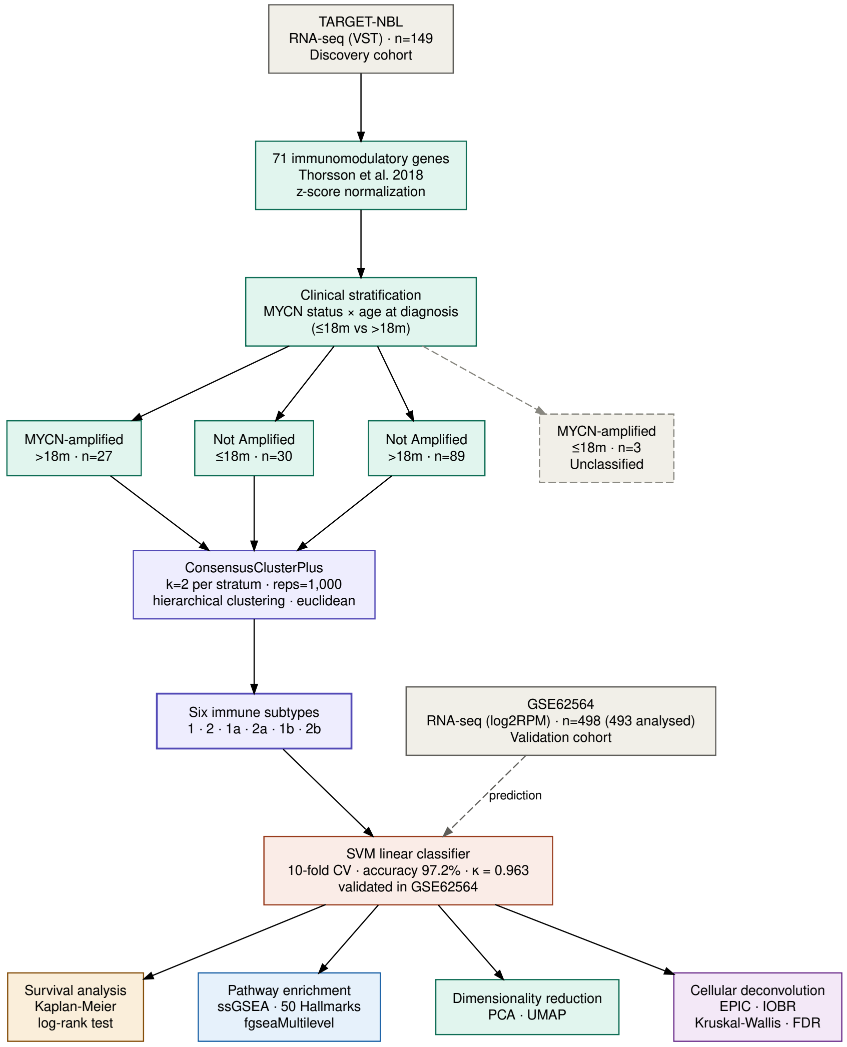


**Fig. S2.** Analytical pipeline overview. Patients from the TARGET-NBL cohort were stratified by MYCN amplification status and age at diagnosis, followed by consensus clustering of immunomodulatory gene expression profiles, SVM classifier training, and validation in the GSE62564 cohort. Downstream analyses included survival analysis, pathway enrichment, and cellular deconvolution.

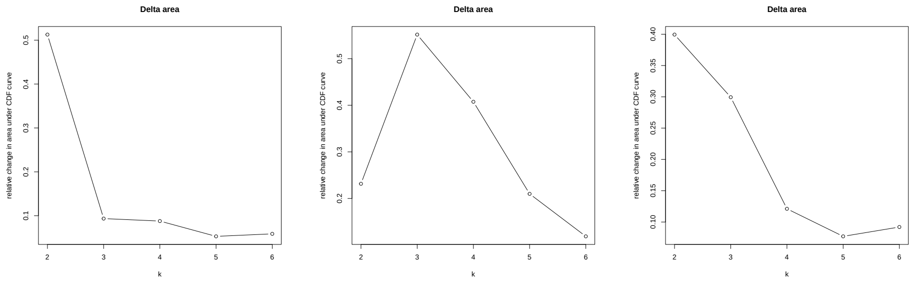


**A** **B** **C**

**Fig. S3.** Delta area plots for consensus cluster number selection in each of the three patient strata: **(A)** MYCN-amplified, age >18 months (n = 27); **(B)** MYCN-non-amplified, age ≤18 months (n = 30); **(C)** MYCN-non-amplified, age >18 months (n = 89). The relative change in area under the consensus CDF curve is shown for $k = 2$ to $k = 6$. In all three groups, $k = 2$ was selected as the optimal partition based on the delta area criterion.

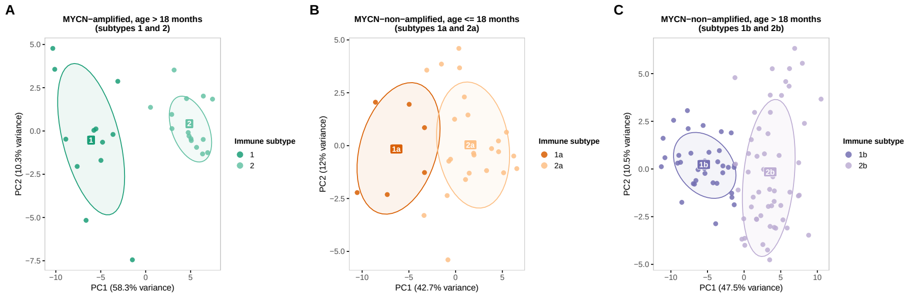


**Fig. S4.** Principal component analysis (PCA) of z-score normalized expression profiles of 71 immunomodulatory genes, performed independently within each clinical stratum. **(A)** MYCN-amplified patients older than 18 months (subtypes 1 and 2; n = 27). **(B)** MYCN-non-amplified patients aged 18 months or younger (subtypes 1a and 2a; n = 30). **(C)** MYCN-non-amplified patients older than 18 months (subtypes 1b and 2b; n = 89). Points are colored by immune subtype; ellipses represent 68% confidence regions per subtype.

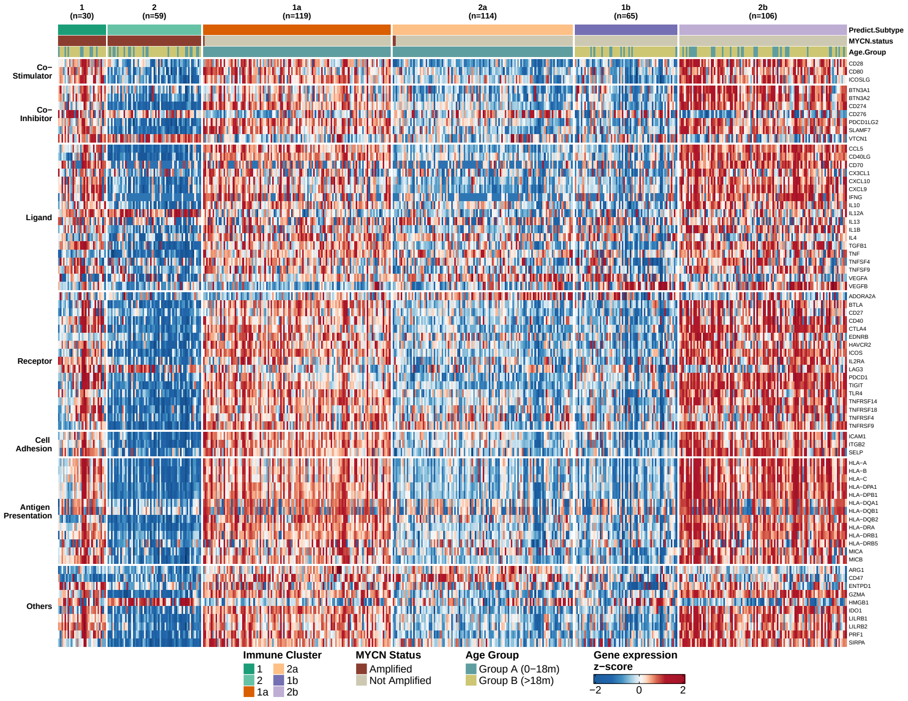


**Fig. S5.** Heatmap of z-score normalized expression profiles of 71 immunomodulatory genes in the GSE62564 validation cohort ($n = 493$), ordered by predicted immune subtype. Gene rows are organized by functional category.

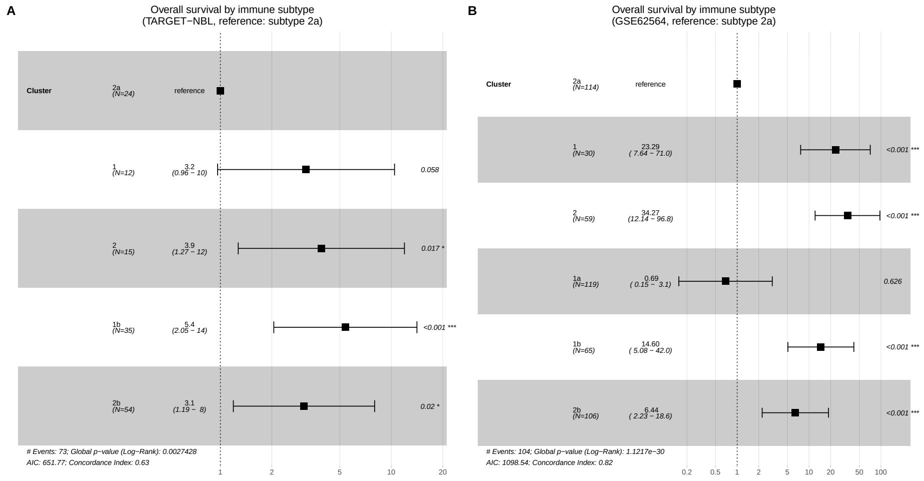


**Fig. S6.** Cox proportional hazards regression for overall survival by immune subtype. **(A)** TARGET-NBL cohort (n = 140, excluding subtype 1a due to complete separation); **(B)** GSE62564 validation cohort (n = 493). Subtype 2a was used as the reference category in both models. Squares represent hazard ratio estimates; horizontal lines represent 95% confidence intervals.

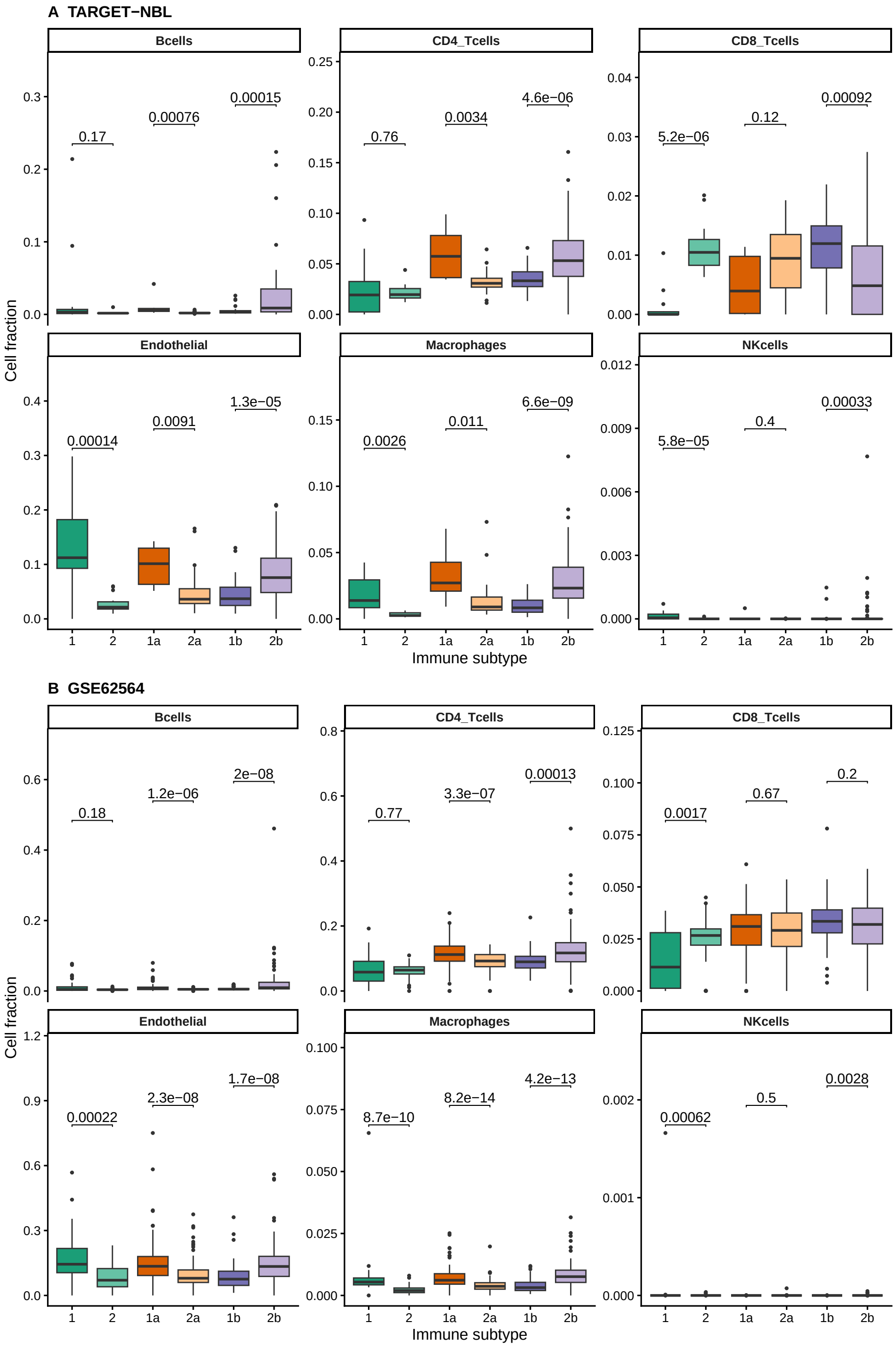


**Fig. S7.** Estimated immune cell fractions by immune subtype. Cellular deconvolution using EPIC was applied independently to the **(A)** TARGET-NBL ($n = 146$) and **(B)** GSE62564 ($n = 493$) cohorts. Boxplots show the distribution of estimated cell fractions for six immune and stromal cell types across the six immune subtypes. Statistical comparisons between paired subtypes within the same clinical stratum (subtypes 1 vs. 2, 1a vs. 2a, and 1b vs. 2b) were performed using pairwise Wilcoxon tests with false discovery rate correction; adjusted p-values are shown above each comparison. Cell fraction estimates reflect proportions of the total cell mixture as estimated by EPIC and should be interpreted as transcriptional proxies of immune cell abundance rather than direct measures of cellular infiltration.
